# Structural basis for antibody-mediated neutralization of Lymphocytic choriomeningitis virus

**DOI:** 10.1101/2022.07.17.500368

**Authors:** Alex Moon-Walker, Zeli Zhang, Dawid S. Zyla, Tierra K. Buck, Haoyang Li, Ruben Diaz Avalos, Sharon L. Schendel, Kathryn M. Hastie, Shane Crotty, Erica Ollmann Saphire

## Abstract

The mammarenavirus Lymphocytic choriomeningitis virus (LCMV) is a globally distributed zoonotic pathogen that can be lethal in immunocompromised patients and cause severe birth defects if acquired during pregnancy. Despite the fundamental importance of LCMV for studying immunobiology, the structure of the trimeric surface glycoprotein, essential for entry, vaccine design and antibody neutralization, remains unknown. In this study, we present the cryoEM structure of the LCMV surface glycoprotein (GP) in its trimeric prefusion assembly both alone and in complex with a rationally engineered monoclonal neutralizing antibody termed 18.5C-M28 (M28). Additionally, we show that passive administration of M28 protects mice from LCMV clone 13 (LCMV^cl13^) challenge when administered as either a prophylactic or therapeutic. Our study illuminates not only the overall structural organization of LCMV GP and the mechanism for its inhibition by M28, but also presents a promising therapeutic candidate to prevent severe or fatal disease in individuals who are at risk of infection by a virus that poses a threat worldwide.

**Highlights:**

- Rationally-engineered antibody M28 neutralizes lymphocytic choriomeningitis virus *in vitro*.
- First high-resolution cryoEM structure of the pre-fusion trimeric lymphocytic choriomeningitis virus glycoprotein alone and in complex with M28.
- M28 neutralizes by bridging adjacent glycoprotein protomers and locking it in the pre-fusion state.
- Prophylactic and therapeutic administration of M28 protects mice from chronic lymphocytic choriomeningitis virus infection.

## Introduction

The prototypic mammarenavirus Lymphocytic choriomeningitis virus (LCMV) is a zoonotic pathogen transmitted by mice that frequent human habitations. LCMV is endemic in every populated continent due to the global distribution of its rodent host, and has served as a critical model to study virus-host interactions since its discovery in 1934 ^1–7^. Although infection of healthy individuals presents with minimal symptoms or symptoms not requiring medical care, LCMV is known to cause severe disease and fatalities in a subset of individuals. For example, LCMV can be lethal if acquired by immunocompromised individuals, such as transplant patients, and can cause birth defects and fetal demise if acquired during pregnancy^8–12^. Furthermore, laboratory exposures to LCMV from working with infected animals as well as from accidental needle sticks have been documented^13–15^.

The glycoprotein (GP) of LCMV is the sole protein on the surface of virions. GP mediates cell attachment and entry, and is the primary target for neutralizing antibodies. In producer cells, the glycoprotein precursor GPC is inserted into the endoplasmic reticulum (ER) membrane, where host signal peptidase (SPase) cleaves the N-terminal stable signal peptide (SSP), resulting in a non-covalently associated complex of SSP and GP (Fig. S1A)^16, 17^. Unlike most signal peptides, the arenavirus SSP remains associated with GP and plays critical roles in GP trafficking, processing, and pH sensing^16–19^. After SPase cleavage of the SSP, the SSP-GP complex is trafficked to the Golgi compartment, where GP is recognized by the host site 1 protease (S1P) and is cleaved into the receptor-binding GP1 and the fusion-mediating GP2 subunits^19–23^. Mature GP assembles on the cell surface as a trimer, with each monomer composed of non-covalently associated SSP, GP1 and GP2 subunits. During virus entry, LCMV GP first engages matriglycosylated ɑ-Dystroglycan (ɑ-DG) and heparan sulfate^24–29^. After viral uptake to acidic endo-lysosomal compartments, LCMV GP undergoes a receptor switch to sialomucin CD164^29^. Conformational changes initiated by CD164 engagement ultimately lead to GP2-mediated membrane fusion^29–31^.

Monoclonal antibodies from patients infected with LCMV have not yet been identified. Antibodies raised in mice typically have lower affinity and are non-neutralizing^32^. Some antibodies isolated from mice can reduce viral load *in vivo* at late stages of infection by enhancing the inflammatory pathway through activation of the cytotoxic T lymphocyte (CTL) response^33–37^. The few neutralizing antibodies against LCMV that have been identified target the GP1 subunit alone, but the precise epitopes have not yet been elucidated. These antibodies confer protection in both wild-type (C57BL/6) mice and in a CTL-deficient murine model, but neutralization-resistant virus mutants are selected in the absence of CTL^35^. Studies of the immune response against Lassa virus (LASV), a close relative of LCMV and etiological agent of Lassa hemorrhagic fever, revealed that most neutralizing antibodies elicited during Lassa virus infection target the pre-fusion conformation of GP^38–42^. These antibodies bind to four discrete epitopes on the surface of GP^38–40, 43, 44^, although the majority target the epitope termed GPC-B. GPC-B antibodies bind LASV GP at the base of the trimer and lock GP in the pre-fusion conformation by engaging adjacent GP monomers^40^. Notably, some GPC-B antibodies cross-react with LCMV GP^38^, suggesting that they could be engineered to neutralize LCMV infection. Since poor neutralizing antibody responses to LCMV are mounted by virus-inoculated mice and there are no therapeutic interventions against this global pathogen, engineering and/or discovery of therapeutic neutralizing antibodies against LCMV is critical.

Here we present cryo-EM structures of the pre-fusion LCMV GP trimer alone and in complex with a LASV-elicited GPC-B antibody that is engineered to neutralize LCMV. We further demonstrate that this human antibody is protective against LCMV infection, in both prophylactic and post-exposure modes. These studies illuminate new features of the LCMV GP trimer and provide a basis for the development of therapies to treat LCMV infections.

## Results

### Engineering a pre-fusion LCMV GP trimer

The architecture of the processed, prefusion glycoprotein trimer that recapitulates the authentic GP presented on the surface of LCMV virions has yet to be elucidated, although the structure of an uncleaved LCMV GP monomeric precursor is known^45^. To date, the only trimeric prefusion arenaviral glycoprotein that has a high-resolution structure is LASV GP, which was solved using a combination of antibodies and site-directed mutations to stabilize the trimeric assembly and maintain the metastable trimer in a pre-fusion state^39, 40^. A recent structure of full-length (membrane-containing) LASV GPC revealed that the engineered, pre-fusion ectodomain accurately mimics the conformation present at the plasma membrane and on the virion surface^40, 46, 47^. We built on this successful engineering strategy for structure determination of LASV GP to engineer and produce a stable, pre-fusion LCMV GP trimer, which we term LCMV pfGP-TD (Fig. S1A). This stabilized construct contains: (a) di-cysteine mutations G207C and G366C in GP1 and GP2, respectively, to ‘staple’ the non-covalently associated subunits together; (b) a E334P mutation in the hinge region within the HR1 of GP2 to prevent GP from rearranging to the lower-energy post-fusion state; (c) replacement of the native S1P cleavage site (RRLA) with that of furin (RRRR) to enhance processing in insect producer cells; (d) an S398A mutation to remove the *N*-linked glycan site at N396 that sterically hinders access to the GPC-B epitope^39^ and (e) an exogenous trimerization domain^48^ at the GP2 C terminus that promotes trimerization (Fig S1A). Biochemical analysis and negative stain microscopy of this construct revealed that the majority of the purified engineered protein was fully processed and trimeric (Fig. S1B-C).

### Monoclonal antibody M28 potently neutralizes LCMV *in vitro*

Some monoclonal antibodies (mAbs) from human survivors of Lassa fever demonstrate cross-reactivity to LCMV GP^38, 40^. Most of these cross-reactive antibodies belong to the GPC-B competition group, which comprises an epitope that bridges neighboring GP monomers together at the base of the GP trimer. We first examined the ability of the GPC-B mAbs 37.7H and 18.5C to neutralize authentic LCMV *in vitro*. Although 37.7H can recognize LCMV GP present at the cell surface^38^, it does not neutralize LCMV (Fig. 1A). In contrast, mAb 18.5C^39^ had moderate neutralization activity toward LCMV with an IC_50_ of 3.77 µg/mL.

**Figure 1.**
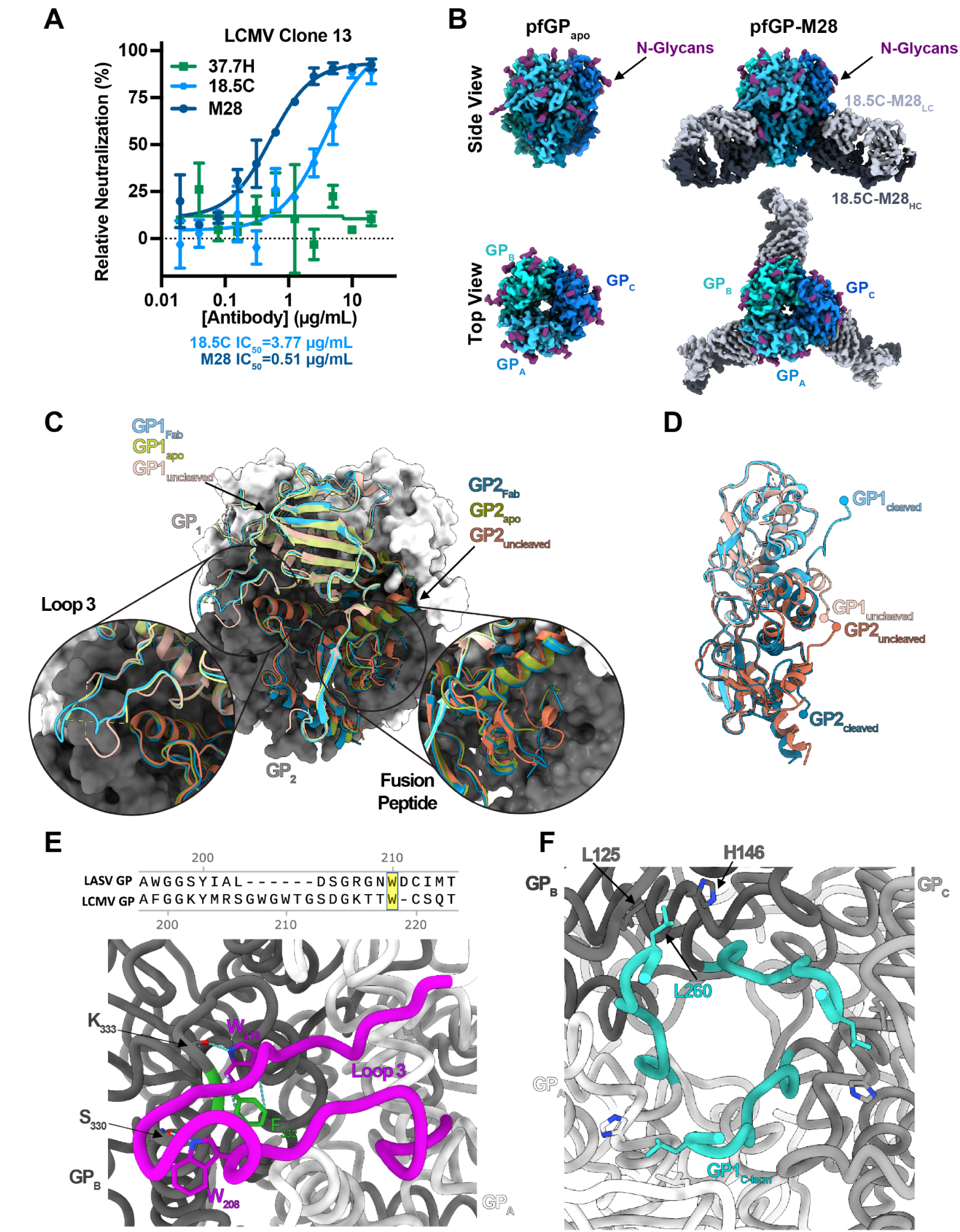
Organization of the pre-fusion LCMV GP trimer. A) The neutralization activity of GPC-B mAbs 37.7H, 18.5C and M28 against authentic LCMV^cl13^ tested in an immunofocus assay with Vero cells. Infectivity data were normalized to control wells containing no antibody. Error bars indicate the standard deviation of the mean from two independent experiments, each performed in duplicate. B) Side and top views of cryoEM density maps of LCMV pfGP-TDapo (left) and the LCMV pfGP-TD-M28 complex (right). Each GP monomer is colored in a different shade of blue, with the lighter and darker shades denoting GP1 and GP2, respectively. The portion of M28 Fab with observable EM density is colored light gray (light chain) and dark gray (heavy chain). N-linked glycans are colored purple. C) Comparison of an uncleaved protomer from previously determined the LCMV GP monomer crystal structure (“GP_uncleaved_”, PDB 5INE) with a cleaved protomer from the cryoEM structures of LCMV pfGP-TDapo (“GP_apo_”; light and dark green for the GP1 and GP2 subunits, respectively) and LCMV pfGP-TD-M28 (“GP_Fab_”; light and dark blue for the GP1 and GP2 subunits, respectively). GP monomers B and C are shown in surface representation, while monomer A is shown in cartoon representation. The uncleaved crystal structure of the GP monomer (light and dark orange for the GP1 and GP2 subunits, respectively) and the cleaved pfGP-TDapo structure were aligned to GP monomer A from the antibody-bound LCMV pfGP-TD. Circled insets highlight trimeric interface interactions mediated by Loop 3 in GP1 and the variable location of the fusion peptide of LCMV GPapo and GP-M28 as compared to the uncleaved LCMV GP monomer. D) Superimposition of uncleaved and cleaved (pfGP-TD-M28) LCMV GP monomers. The C terminus and GP1 and N terminus of GP2 for both structures are indicated with spheres. E) Sequence alignment of loop 3 within LCMV GP with corresponding residues in Lassa virus GP shown at the top. In the fully cleaved/processed LCMV trimer, loop 3 in GP1 forms a pi-pi stacking interaction and hydrogen bonds with a neighboring GP monomer. GP monomers A and B are colored light and dark gray, respectively. Loop 3 in monomer B (encompassing residues 199-224) is colored magenta, with GP monomer A residue F332 shown in green. The GP monomer B residues W208 and W219 are shown in magenta. F) Apical stabilization of the trimeric interface of GP. The C-terminal loop of GP1 is colored cyan with the major matriglycan recognition determinant L260 residue shown with its interacting partners L125 and H146.

We also tested 18.5C-M28 (M28), which was rationally engineered from 18.5C to be more potent against different LASV lineages^39^. M28 incorporates an S32R mutation in the CDR H1 and has an insertion of R55 in CDR H2 (R55ins) (Fig. 1A). Similar to the results for LASV, M28 could neutralize LCMV with 7.4-fold greater potency over parental 18.5C^39^ (Fig. 1A).

### Structural organization of the LCMV GP trimer

Antibody M28 binds a conformationally dependent, pre-fusion specific epitope and readily recognizes recombinant LCMV pfGP-TD (Fig. S1D), indicating that our soluble construct is in the pre-fusion conformation. Furthermore, analysis of the LCMV pfGP-TD-M28 Fab complex by negative stain EM showed that the engineered GP was bound by three M28 Fabs (Fig. S1E-G).

To elucidate the structure of the pre-fusion LCMV GP trimer and understand if conformational changes occurred upon antibody binding, we obtained three-dimensional cryoEM reconstructions of the engineered trimer alone (pfGP-TDapo) and in complex with M28 Fab (pfGP-TD-M28), both at 3.2 Å resolution (Fig. 1B; Table S1; Fig. S2A-F, Fig. S3A-E). Comparing the structure of the GP monomers in this pre-fusion cleaved trimer to that determined for the uncleaved LCMV GP monomeric protomer^45^, shows that the overall architecture of the GP is essentially the same. However, there are three differences. First, loop 3^45^, encompassing residues 207-219 of GP1, is now fully resolved in the context of the trimer in complex with M28 (Fig. 1C, Fig. S3A). This loop lies at the trimeric interface and reaches out from one protomer to interact with the neighboring protomer. In particular, residue W208 of GP1-A forms a hydrogen bond with the backbone of GP2-B residue 330 in HR1, while residue W219 of GP1-A forms a pi-stacking interaction with residue F332 and hydrogen bonds with the backbone-oxygen of K333 of GP2-B (Fig. 1E, Fig. S3C). This interaction is similar to that observed for the equivalent tryptophan (W210) in LASV GP^44, 46^.

A second difference between uncleaved GP and the cleaved GP trimer structures described here, is a ∼25 Å and ∼20 Å displacement of the C terminus of GP1 and N terminus of GP2, respectively, from their position in the uncleaved structure (Fig. 1D). The new termini are created by S1P cleavage of the GP1-2 protomer^20–22^. The large translation of the C terminus supports the required role of S1P cleavage for GP trimer formation: the C and N termini of GP1 and GP2, respectively, must separate following cleavage to allow proper trimer formation, as was similarly noted for LASV^40, 46^. Transposition of the GP1 C terminus also allows formation of an additional stabilizing interaction with the neighboring GP monomer, whereby residue L260 of monomer A interacts with L125 and H146 of monomer B (Fig. 1F). Residue 260 has been shown to be a major determinant for receptor binding across different LCMV strains, with the leucine of GP from Clone 13 and WE-HPI strains conferring high affinity and the phenylalanine of Armstrong strain GP conferring low affinity (Fig. 1F, Fig. S3E)^25, 49, 50^. Interaction of the C terminus of GP1 with neighboring monomers is also critical for trimer formation and matriglycan recognition for Lassa virus^40, 45, 46, 51, 52^, suggesting that matriglycan-dependent utilizing arenaviruses share a conserved mechanism for trimer stabilization and receptor recognition.

Lastly, we demonstrated that the fusion peptide (266-272) present at the N terminus of the GP2 subunit and freed from GP1 by cleavage of the GP1-2 precursor, has a different location than that predicted based on the Lassa GP trimer. The N termini of the fusion peptides of Lassa GP trimers solved to date are each situated at the trimeric interface and are in a near-identical location regardless of whether or not an antibody is bound (Fig. 2A)^39, 40, 43, 44, 46^. Modeling of one protomer of uncleaved LCMV GP onto the Lassa GP trimer predicted the GP1 C-terminal transposition, but did not necessarily suggest that any more than local repositioning of the GP2 N-terminus occurred. However, we find that LCMV fusion peptide is more flexible than that of LASV GP and rather than gathering at the center of the trimer, the LCMV fusion peptide appears to be oriented in a more outward presentation. Indeed, in both the apo and M28-bound LCMV GP structures, only residues W270-S273 can be modeled. These residues interact with HR1a within the same protomer and the HR1c of another to form a ∼90° path rather than tracking straight between HR1b and HR1c of two adjacent monomers and into the center of the trimer (Fig. 2A; Fig. S3B).

**Figure 2.**
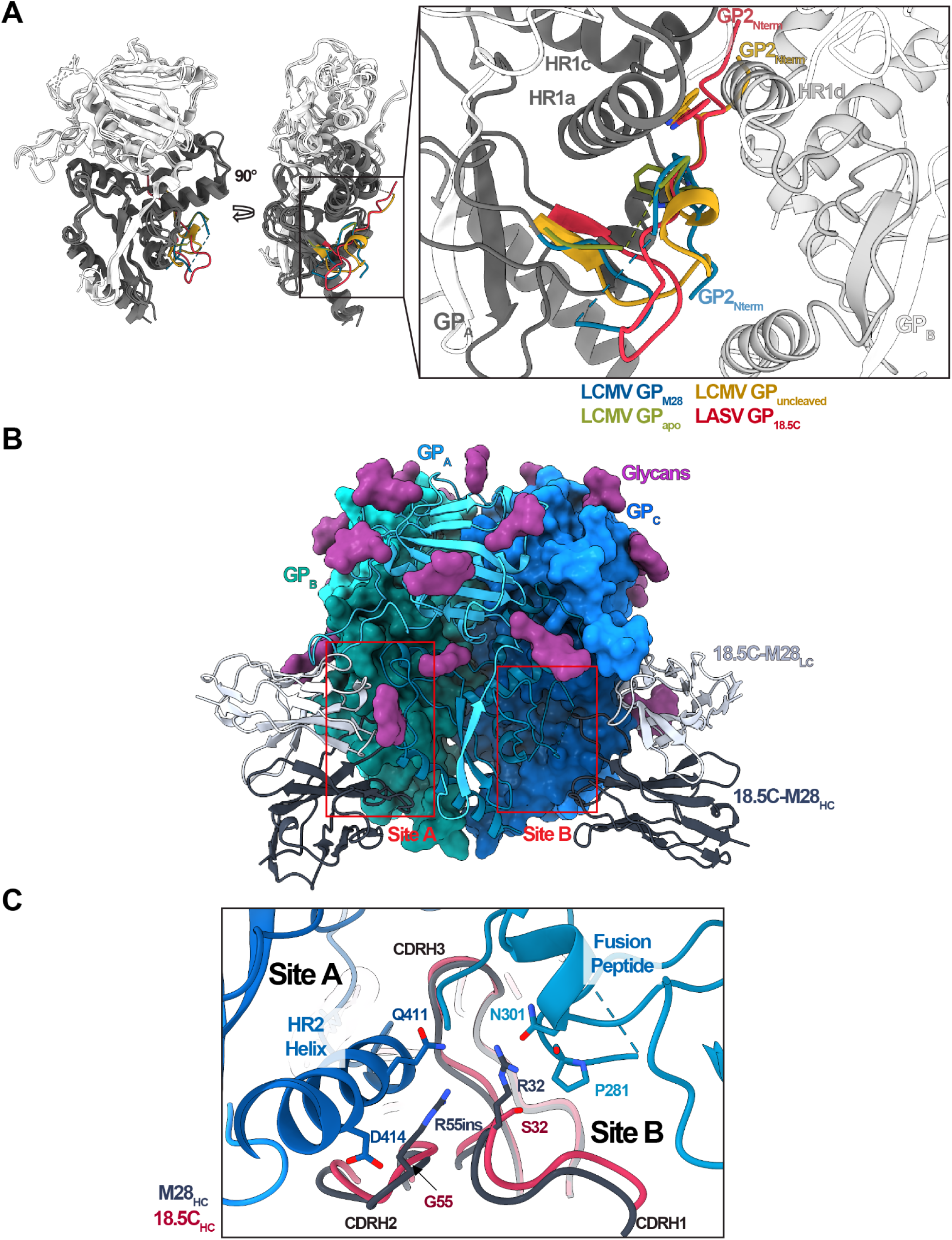
Recognition of LCMV GP by monoclonal antibody M28. A) Labile conformation state of the fusion loop shown in front (left) and side (middle) views. The structures of monomeric Lassa and LCMV GP are overlaid and colored in light and dark gray to depict GP1 (white) and GP2 (dark gray) subunits, respectively. Lassa pfGP residues 260-279 (PDB:6P91), uncleaved LCMV pfGP residues 267-285 (PDB:5INE), cleaved LCMV pfGP (pfGP-M28: residues 267-285), and cleaved LCMV pfGPapo (residues 270-285) are colored red, yellow, blue and green, respectively. B) GP monomers B and C are shown in surface representation, while monomer A is shown in cartoon representation. M28 light and heavy chains are shown in light and dark gray, respectively. Recognition sites A and B are highlighted by red boxes. C) The complementarity determining regions (CDRs) of the light (light gray) and heavy (dark gray) chains of M28 are shown in cartoon representation. M28 residues R55 and R32 are shown in stick form in binding site A (left) and B (right), respectively. The CDRs of 18.5C (PDB: 6P91) are overlaid and are colored light pink and red for the light and heavy chains, respectively. The parental R55 and S32 residues of 18.5C are shown in stick form. LCMV GP1 and GP2 are colored in light and dark blue, respectively.

### Structural organization of the LCMV GP trimer and its recognition by mAb M28

The structure of the LCMV GP-M28 Fab complex reveals that M28 recognizes a quaternary footprint that involves simultaneous contact with two neighboring GP monomers (Fig. 2B). M28 binding buries ∼1,457Å^2^ of its molecular surface overall, similar to the surface area that 18.5C engages on Lassa virus GP. Like other antibodies in the GPC-B competition group, M28 binding bridges two monomers together in a bipartite recognition involving site A, which encompasses primarily the HR2 region of monomer A, and site B, which encompasses primarily the fusion peptide on the adjacent monomer B. Site A has the majority of the surface area buried in the M28 interaction, and the amount buried is larger than that seen for the parental 18.5C on LASV site A (1,237 Å^2^ vs. 935Å^2^). Concomitantly, M28 recognizes a smaller area on LCMV site B than parental M28 does on LASV (220 Å^2^ and 580 Å^2^ for LCMV and LASV GP, respectively).

The rationally engineered antibody M28, bears a S32R mutation in CDR H1 and an arginine insertion at residue 55 (R55) in CDR H2, which together contribute to increased in potency against all Lassa virus lineages compared to parental 18.5C^39^. Here, we similarly find that M28 has a 7.4-fold higher potency against LCMV compared to 18.5C (Fig. 1A). Based on an alignment of Lassa virus and LCMV GP structures, we hypothesized that M28 residues R32 and R55 would interact with site A residues Q411 and D414, respectively, within HR2 of LCMV GP. While some of the aliphatic portion of R32 is ordered, we cannot see density for the guanidinium group (Fig. S3D), which may interact with site B through the backbone oxygen of P281 in the fusion peptide and N301 in the fusion loop (Fig. 2C; Fig. S3D). R55 has clear side-chain density (Fig S3D), and is positioned ∼180° from D414 and towards Q411 (Fig. 2C; Fig. S3D). Superimposition of the crystal structure of 18.5C^39^ over that of M28 suggests that, just like in the Lassa GP-18.5C complex^39^, the parental 18.5C residues S32 and G55 would have insufficient length to interact with GP at either site A or B (Fig. 2C).

In the LCMV pfGP-M28 complex structure, loop 3 hugs and sits at the top of the M28 light chain (Fig. 1C). We previously identified this region as modulating access by GPC-B antibodies to Lassa Lineage I GP^43^. LCMV, along with its close relatives Lunk virus^53^ and Dandenong virus^54^, have the longest loop 3 amongst arenaviruses. The pfGPapo structure has incomplete density for loop 3 (Fig. 1C, Fig. S3A), indicating that, in the absence of the antibody, loop 3 is flexible and might partially occlude the GPC-B epitope. This mechanism of GPC-B antibody evasion could explain the lack of neutralization by 37.7H, as observed with Lassa Lineage I^43^. Furthermore, to improve neutralizing responses and to elicit neutralizing antibodies targeting the GPC-B epitope, partial or complete removal of loop 3 might be necessary .

Due to stabilizing interactions between M28 and LCMV GP, we could model residues T267-D274 and D280 onward with W270-S273 packing in the same conformation as in pfGPapo (Fig. 2A). The lack of density in the apo structure compared to the antibody-bound structure may reflect this region’s lability, yet the observable and identical conformation of residues W270-S273 in both structures suggests that the fusion peptide conformation is an inherent feature of LCMV GP and not conferred by antibody binding. The adoption of such different conformations between the fusion loops from Lassa and LCMV when bound by neutralizing GPC-B antibodies suggests that neutralization depends primarily on site A interactions for neutralization. As such, further rational mAb design should focus on improving site A recognition for increased potency and cross-reactivity to other arenaviruses.

### Passive administration of M28 as either a prophylactic or therapeutic protects mice from LCMV^cl13^ challenge

Antibodies raised during the humoral response to LCMV infection in mice are typically non-neutralizing^32, 35, 55^. Those that do neutralize are raised at late points of infection and only provide partial protection^32, 35, 55–58^. Due to the neutralization potency of M28 *in vitro*, we sought to examine its ability to protect mice from LCMV-mediated disease *in vivo*. We first assessed the prophylactic potency of M28 in C57BL/6 mice infected with LCMV^cl13^ (Fig. 3A). Mice were treated with 300 µg (150 µg I.V. and 150 µg I.P.) of either M28 IgG or a control IgG, and one day later, infected with 0.3×10^6^ FFU LCMV^cl13^. Seven days post-infection, M28-treated mice had a 20-fold lower serum viral titer compared to the control IgG-treated mice (Fig. 3B). On day 14 and day 21, no virus was detected in serum from any of the M28-treated mice (Fig. 3B). In contrast, in control IgG-treated mice, virus titers on days 14 and 21 were similar to those on day 7 (Fig. 3B). In tissues, virus was undetectable in spleen, liver and kidneys of M28-treated mice on day 21 (Fig. 3C) and these mice had no infection-induced weight loss compared to control IgG-mice (Fig. 3D). In addition, administration of M28 one day before a high-dose (2×10^6^ FFU) LCMV^cl13^ infection significantly reduced the viral titers in organs and blood on day 3 and day 7 (Fig. S4B-D). On day 2, a 2-fold reduction in viral load in the spleen was observed in the M28-treated mice compared to the IgG control-treated mice (Fig. S4C). On day 7, a 22-, 57-, 9,128-, and 1,961-fold decrease in viral load was seen in the blood, spleen, liver, and kidney, respectively, of M28-treated mice relative to IgG control-treated mice (Fig. S4D).

**Figure 3.**
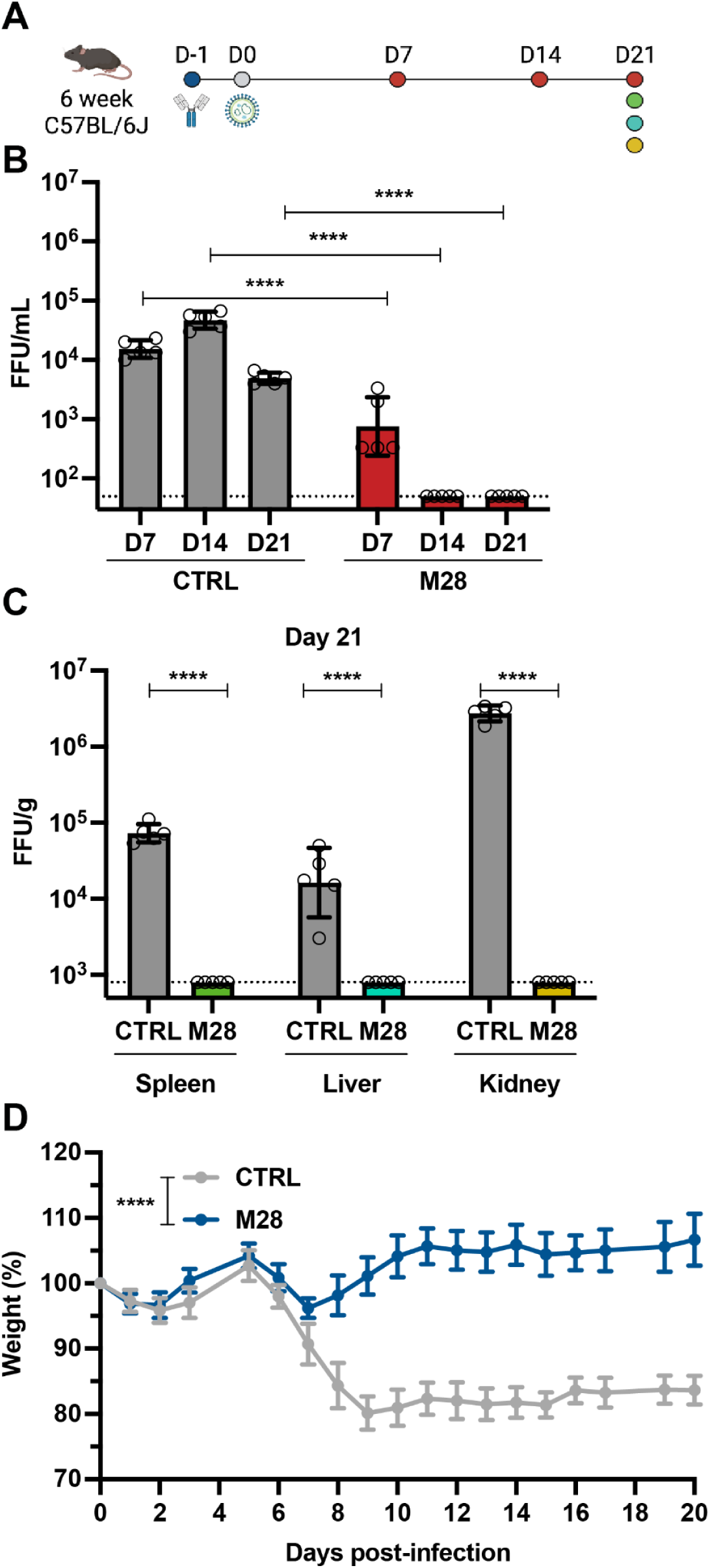
Prophylactic treatment of M28 protected mice from LCMV^cl13^ challenge. A) Schematic of prophylactic M28 administration. M28 or IgG control (CTRL) were administered via I.V. and I.P. on day -1. Mice were infected with a medium dose of LCMV^cl13^ (0.3×10^6^ FFU) via I.V. on day 0. Spleen was harvested on day 3. Blood, spleen, a portion of the liver, and kidneys were harvested on day 7. The organs were subsequently weighed and homogenized for downstream analysis. B) The LCMV viral titers in serum (FFU/mL) on day 7, day 14, and day 21. Two-way ANOVA followed by Tukey’s post hoc test using log-transformed data was used to examine differences between groups. C) LCMV viral titers (FFU/g) in the spleen, liver, and kidneys on day 21. Two-way ANOVA followed by Tukey’s post hoc test using log-transformed data was used to examine differences between groups. D) Mouse body weight was monitored over 20 days. Statistical comparison was performed using two-way ANOVA.

Treatment of disease after establishment of viral infection is a likely clinical scenario. Hence, we also assessed the therapeutic potency of M28 in C57BL/6J mice after infection with LCMV^cl13^ (Fig. 4A). Dose-finding studies indicated that 0.3×10^6^ and 2×10^6^ FFU LCMV^cl13^ infection generated similar viral titers in mouse serum from day 7 to day 28 (Fig. S4A), therefore a 0.3×10^6^ FFU dose of LCMV^cl13^ was used for subsequent studies. Mice were administered 300 µg (150 µg I.V. and 150 µg I.P.) of either M28 IgG or an isotype-matched control IgG two days post-infection (Fig. 4B). Seven days post-infection, M28-treated mice had a serum viral titer of ∼4×10^2^ FFU/mL, while control IgG-treated mice exhibited a 50-fold higher serum viral titer of ∼2×10^4^ FFU/mL (Fig. 4B). Strikingly, on day 14 and day 21, no virus was detected in serum from any of the M28-treated mice (Fig. 4B). In contrast, all animals in the control IgG-treated group had detectable serum viral loads at day 7 and 21 after infection with ∼2×10^4^ FFU/mL and ∼6.6×10^3^ FFU/mL, respectively, indicating viral persistence (Fig. 4B). In tissues, on day 21, virus was undetectable in spleen, liver and kidneys of M28-treated mice (Fig. 4C) and the M28-treated mice had no infection-induced weight loss compared to control IgG-mice (Fig. 4D). These results suggest that M28 can clear LCMV *in vivo* post-infection and prevent the establishment of LCMV-induced chronic disease. Together, these data suggest that M28 can protect mice in both prophylactic and therapeutic contexts and present this antibody as a promising treatment for controlling viral load in at-risk individuals.

**Figure 4.**
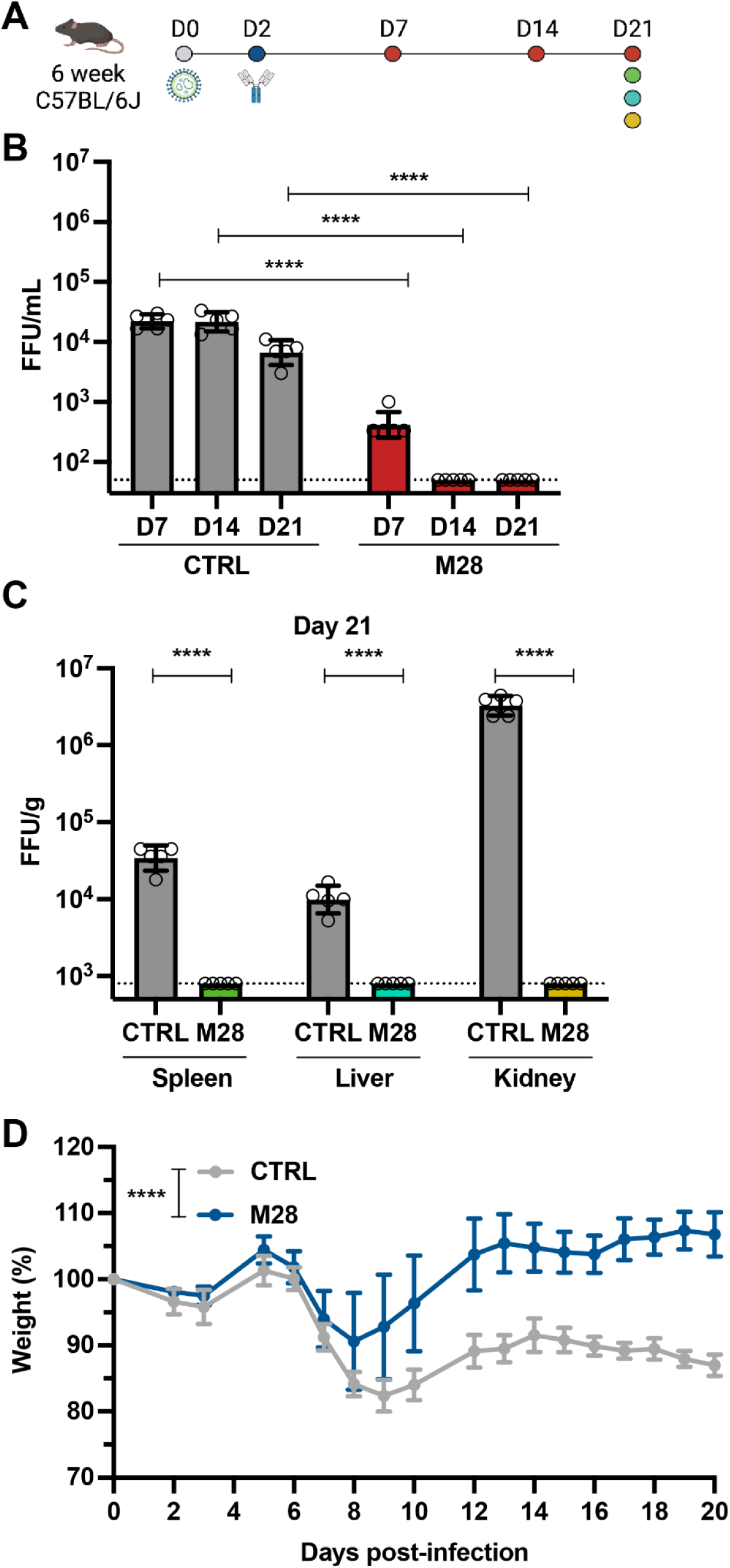
Therapeutic treatment of M28 protects mice from LCMV^cl13^ infection. A) Schematic of therapeutic M28 administration. Mice were infected with a medium-dose (0.3×10^6^ FFU) of LCMV^cl13^ via I.V. on day 0. M28 or IgG control (CTRL) were administered via I.V. and I.P. on day 2. Sera were collected on day 7, 14, and 21. Spleen, a portion of the liver, and kidneys were harvested at day 21, weighed and homogenized for downstream analysis. B) LCMV viral titers in serum (FFU/mL) on day 7, day 14, and day 21. Statistical analysis applied is two-way ANOVA followed by Tukey’s post hoc test using log-transformed data. C) LCMV viral titers (FFU/g) in the spleen, liver, and kidneys on day 21. Statistical analysis applied is two-way ANOVA followed by Tukey’s post hoc test using log-transformed data. D) Mouse body weight was monitored over 20 days. Statistical comparison was performed using two-way ANOVA.

## Discussion

In this study we determined the first high-resolution structure of the trimeric, pre-fusion glycoprotein of LCMV alone and in complex with the rationally-engineered monoclonal antibody M28. We further determined that M28 can neutralize LCMV *in vitro* and protect against disease *in vivo* whether delivered prophylactically or therapeutically. Our trimeric LCMV GP antigen could serve as a basis for improving our understanding of neutralizing responses against LCMV and could potentially be used for both rational vaccine design and antibody therapeutic discovery.

Due to well-documented cases of LCMV infections of transplant patients^9,59, 60^, administration of M28 prior to surgery could be a promising strategy to prevent fatalities caused by LCMV in high-risk settings. Furthermore, identification of M28 as a potent neutralizing antibody and therapeutic against LCMV makes this monoclonal antibody an attractive therapy to combat this global zoonotic pathogen.

## Supplemental Table and Figures

**Table S1.**
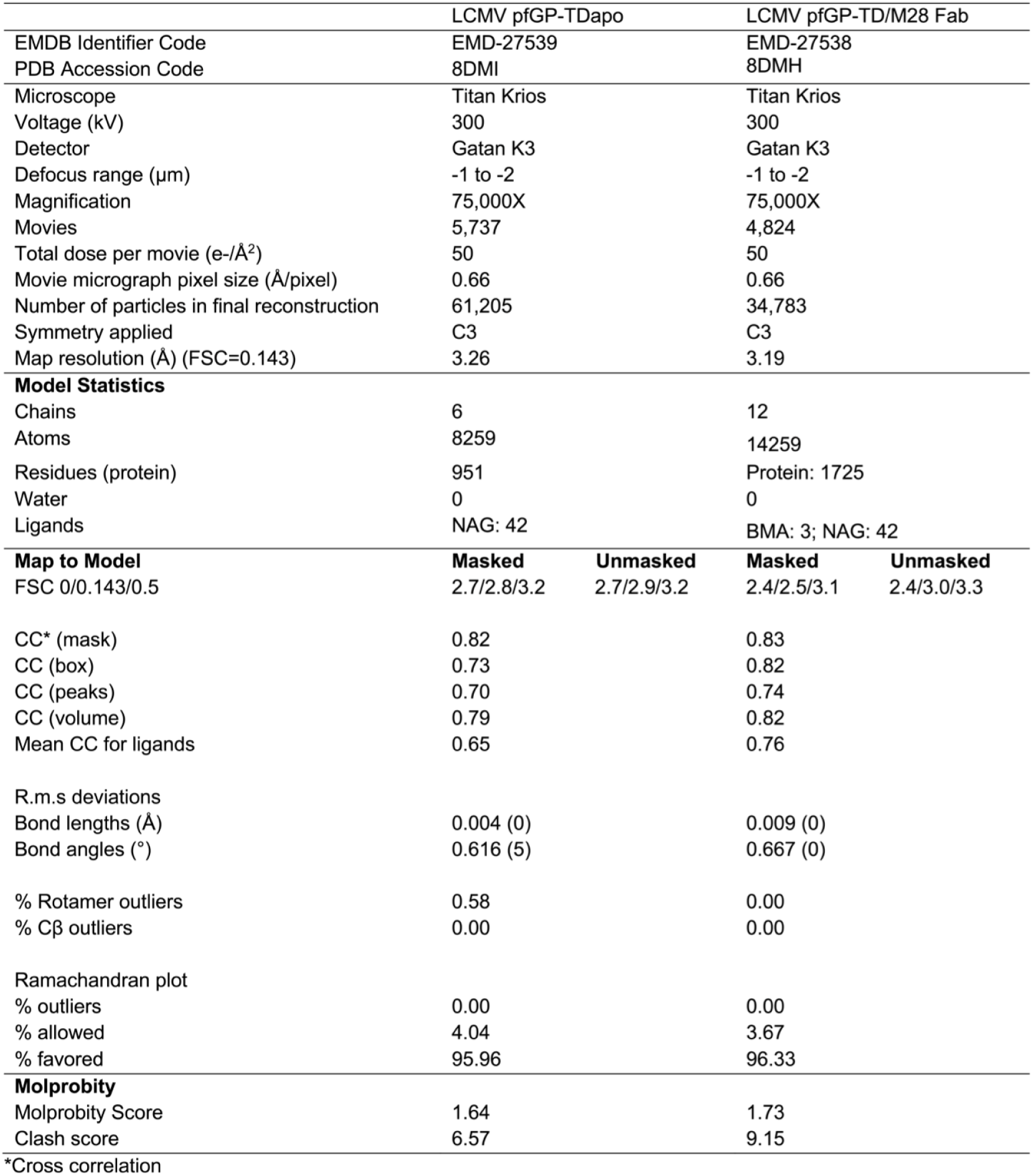
CryoEM map and model statistics for LCMV pfGP-TD alone and in complex with antibody M28; related to Figures 1 and 2.

**Figure S1.**
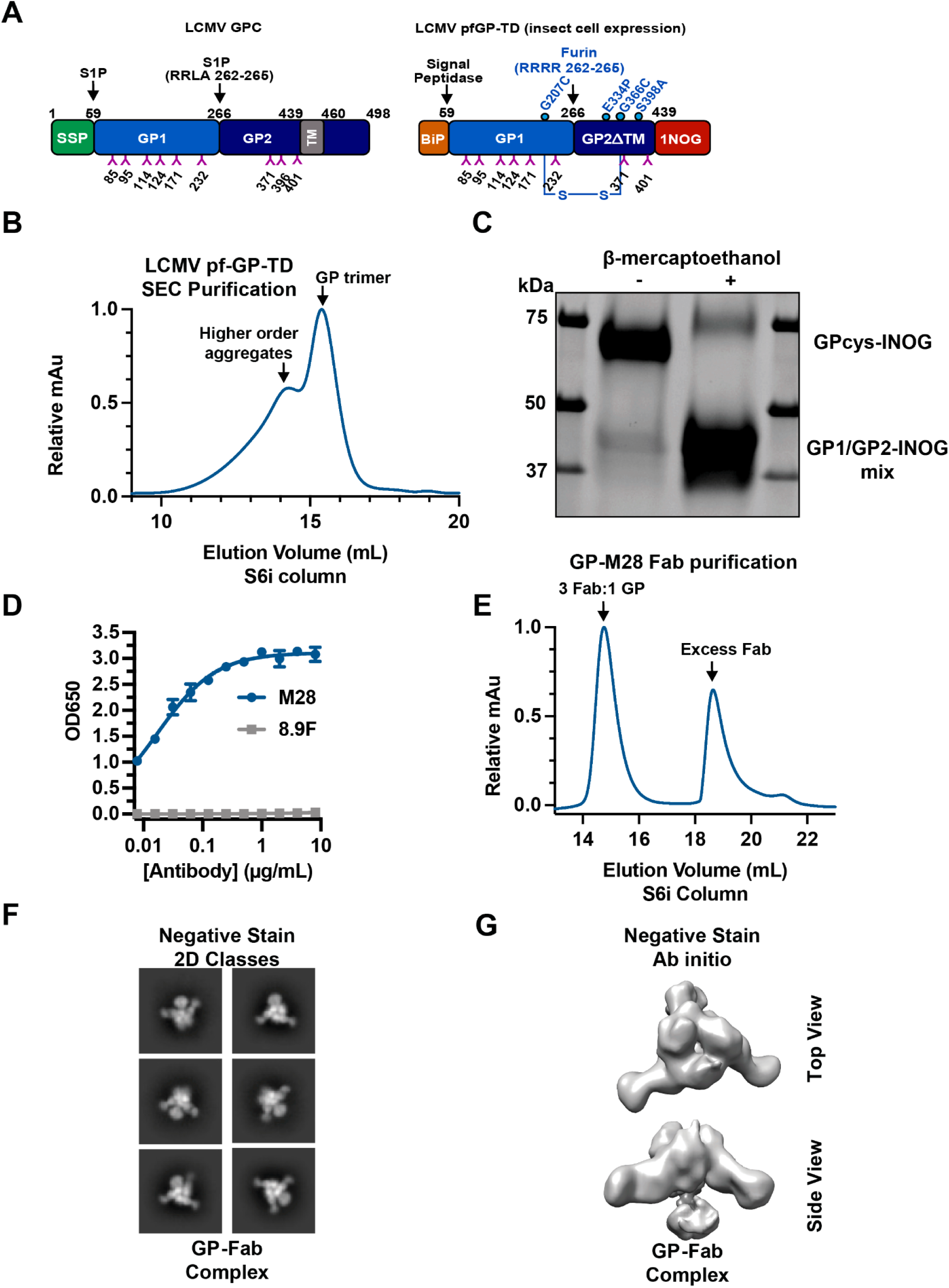
Biochemical and negative stain characterization of the LCMV pfGP-TD-M28 Fab complex; related to Figures 1 and 2. A) Comparison of the topology between full-length LCMV GP (left) and the engineered LCMV pfGP-TD (right). B) Size exclusion chromatography trace of LCMV pfGP-TD separated on a Superose6i column with an arrow denoting the fraction selected for downstream analysis. C) SDS-PAGE analysis of LCMV pfGP-TD under either non-reducing or reducing conditions. D) Soluble LCMV pfGP-TD was coated onto a half-well 96-well, high-protein binding plate. Wells were washed, blocked with 3% BSA, and then incubated with either M28 IgG or 8.9F IgG^38^ (as a negative control) in a 2-fold dilution series for one hour at 37 °C. Wells were washed six times before incubation with goat anti-human IgG-HRP for one hour at 37 °C. Then, the wells were extensively washed and TMB substrate was added, followed by a 650nm stop solution. The plates were then read with a spectrophotometer plate-reader. The experiment was performed twice in duplicate, and a representative ELISA experiment is shown. Error bars indicate the standard deviation of the mean. E) Size exclusion chromatography trace of LCMV pfGP-TD complexed with excess M28 Fab separated on a Superose6i column, with an arrow denoting the fraction selected for downstream negative stain and cryoEM analysis. F) Representative 2D classes of the LCMV pfGP-TD-M28 Fab complex particles analyzed by negative stain EM. G) Ab initio model of the GP/Fab particles obtained from negative stain EM showing the top view (top) and side view (bottom) of the complex.

**Figure S2.**
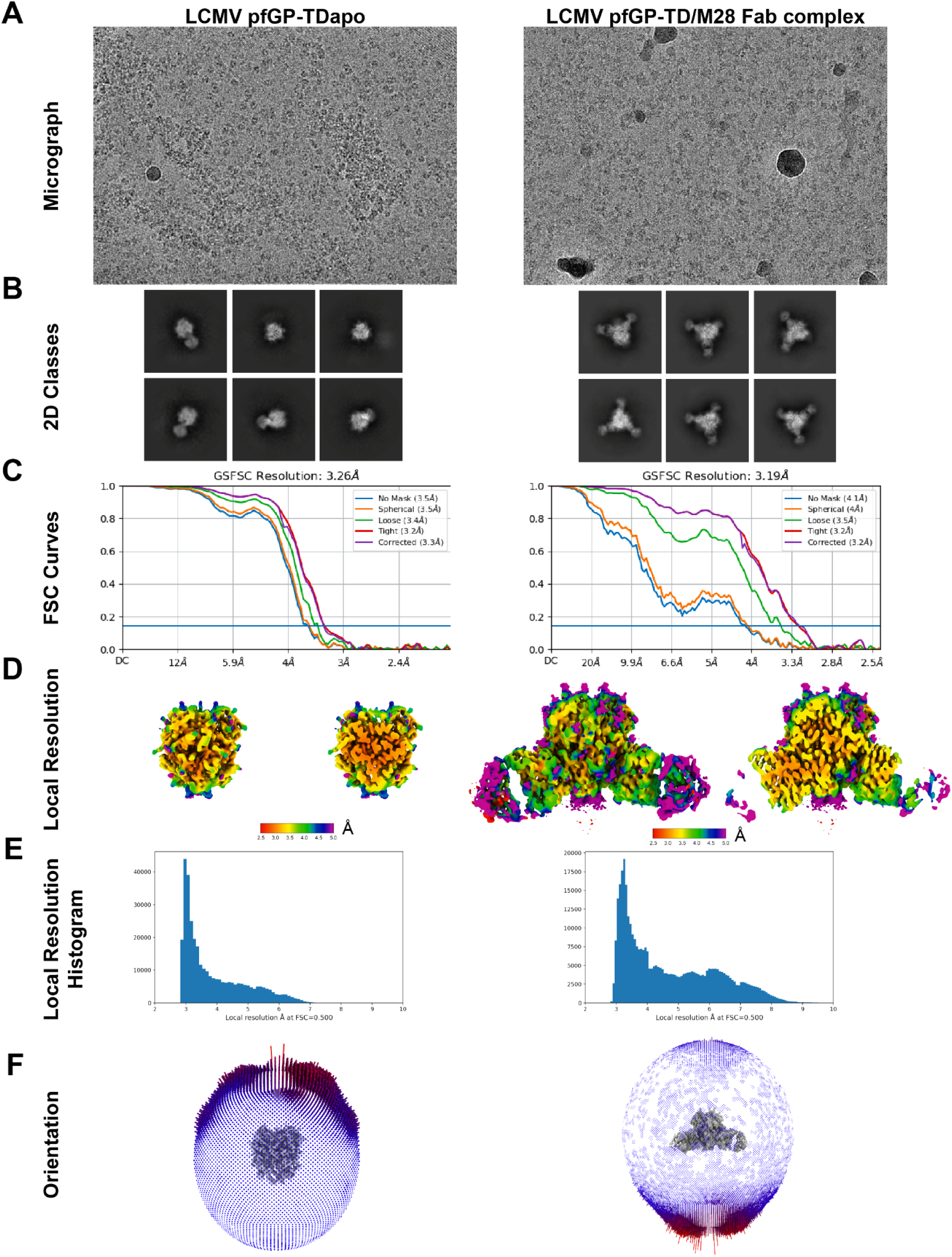
Data processing and validation of LCMV pfGP-TD and pfGP-TD/M28 complex; related to Figures 1 and 2. A) Representative cryoEM micrographs of LCMV pfGP-TDapo (left) and LCMV pfGP-TD/M28 complex (right) particles. B) Representative 2D classes of LCMV pfGP-TDapo (left) and LCMV pfGP-TD/M28 complex (right) particles. C) CryoSPARC Fourier Shell Correlation (FSC) curves of processed cryoEM data sets for LCMV pfGP-TDapo (left) and LCMV pfGP-TD/M28 complex (right). D) Local resolution maps of the LCMV pfGP-TDapo (left) and LCMV pfGP-TD/M28 complex (right). The left volume corresponds to surface resolution while the right volume corresponds to a cross section. E) Local resolution histograms of the LCMV pfGP-TDapo (left) and LCMV pfGP-TD/M28 complex (right) particles. F) Particle orientation distribution for the LCMV pfGP-TDapo (left) and LCMV pfGP-TD/M28 complex (right) cryoEM maps.

**Figure S3.**
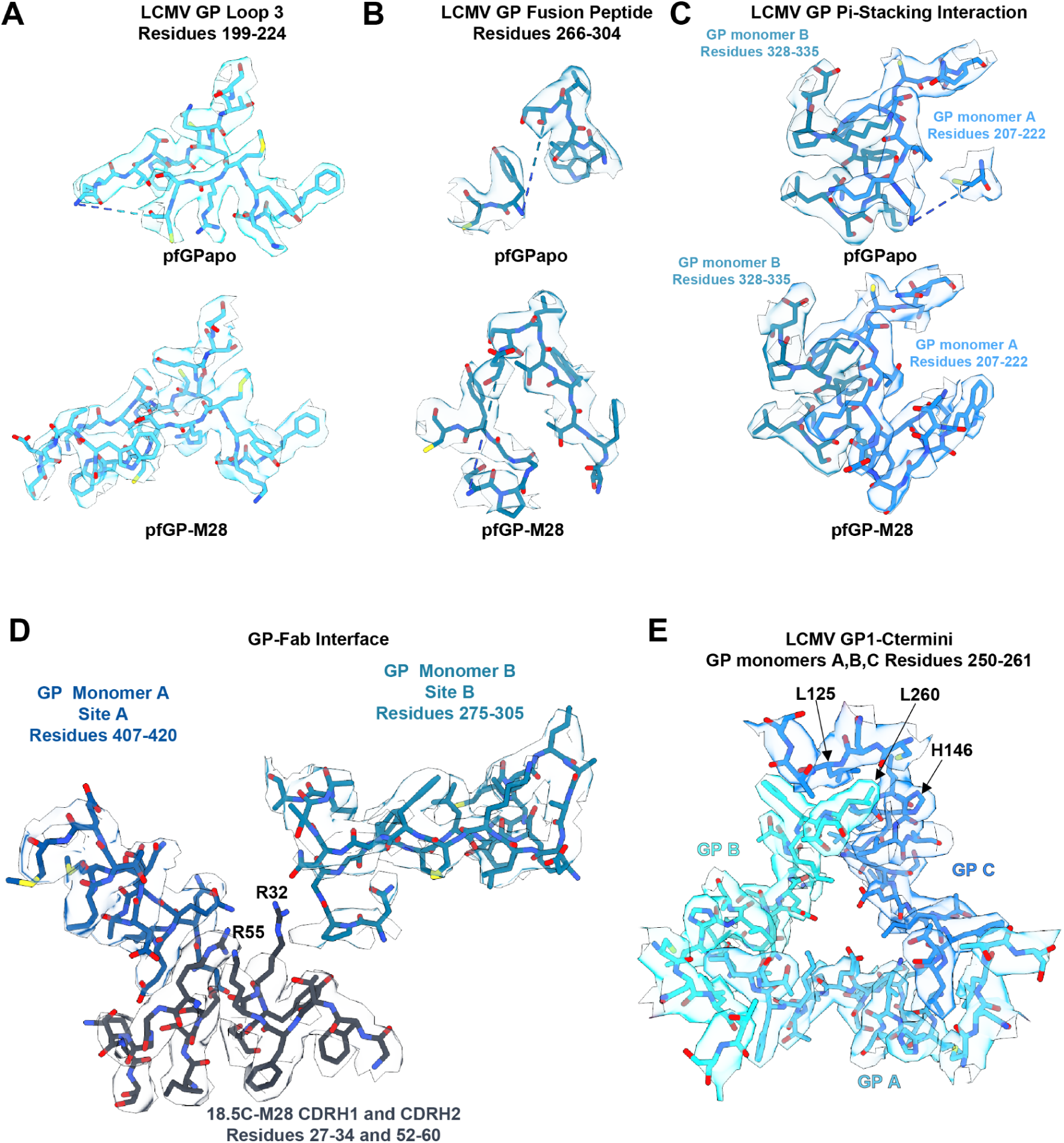
CryoEM density maps with fitted atomic models for areas of interest, related to Figures 1 and 2. A) CryoEM density map of Loop 3 in LCMV pfGP-TD (apo, top; M28-bound, bottom). The cryoEM density map is shown at an isosurface threshold value of 0.1 for Fig. S3A-E. B) CryoEM density map of the fusion peptide (apo, top; M28-bound, bottom) C) CryoEM density map of the pi-stacking interactions between GP protomers (apo, top; M28-bound, bottom). D) CryoEM density map of the GP-Fab interface. In particular, CDR H1 and CDR H2, which have the arginine mutations, are shown along with the fusion peptide at site B and the HR2 helix of site A (apo, top; M28-bound, bottom). E) CryoEM density map of the apical trimeric interface (apo, top; M28-bound, bottom).

**Figure S4.**
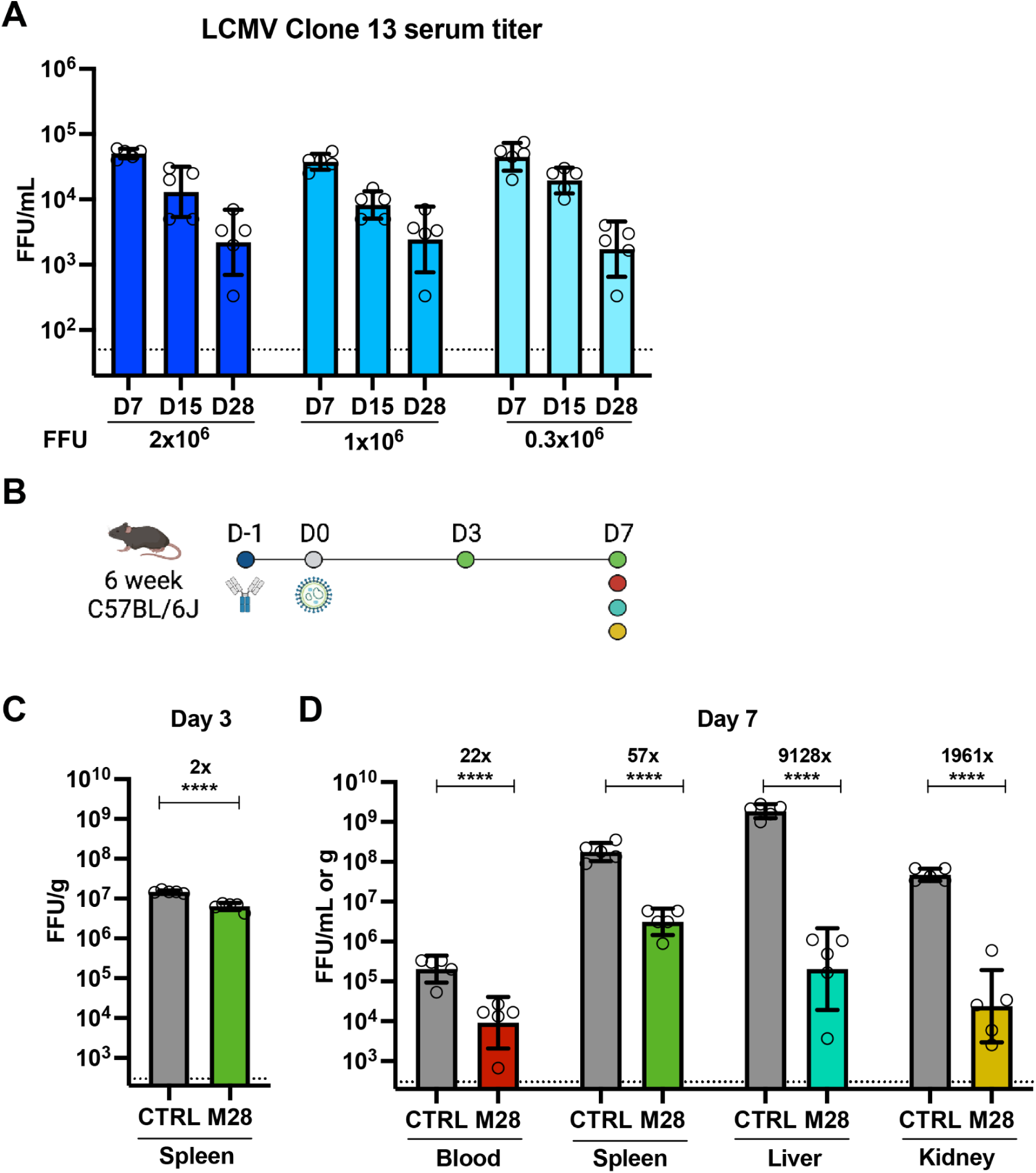
Prophylactic treatment of M28 reduces viral load upon a high-dose LCMV^cl13^ challenge; related to Figure 4. A) Mice were infected with 0.3×10^6^, 1×10^6^, and 2×10^6^ FFU LCMV^cl13^ on day 0. The viral titers in serum were tested on day 7, day 15, and day 28. B) Schematic of prophylactic M28 administration. M28 or IgG control (CTRL) were administered via I.V. and I.P. on day -1. Mice were infected with a high-dose (2×10^6^ FFU) LCMV^cl13^ via I.V. on day 0. Spleens were harvested on day 3. Blood, spleen, a portion of the liver, and kidneys were harvested on day 7. The organs were subsequently weighed and homogenized for downstream analysis. C) LCMV viral titer (FFU/g) in the spleen at day 3. Two-tailed Student’s t-test using log-transformed data was used to assess differences between the groups. Fold-change between control and M28 groups is indicated. D) LCMV viral titers in blood (FFU/mL) and organs (FFU/g) at day 7. Two-way ANOVA followed by Tukey’s post hoc test using log-transformed data was carried out. Fold-changes between control and M28 groups are indicated.

## Acknowledgements

We thank the cryoEM facility of La Jolla Institute for Immunology, philanthropic support of this facility, as well as Dr. Ruben Diaz Avalos for data collection and Drs. Vamseedhar Rayaprolu, Xiaoying Yu and Heather Callaway for their advice on cryoEM processing of the LCMV pfGP-TDapo and LCMV pfGP-TD-M28 complex data. We also thank Nathaniel I. Bloom for helping collect and homogenize mouse organs.

## Funding Sources

EOS: NIH U19 AI 142790; NIH R01 A1132244; NIH R01AI14125 KMH: NIH R21AI137809

AMW: NIH F31-AI154700; NIH T32-AI125179

SC: NIAID P01 AI145815 and LJI Institutional Funds

DZ: Swiss National Science Foundation: P2EZP3_195680, P500PB_210992

## Conflicts of Interest

None.

## Key Resources Table

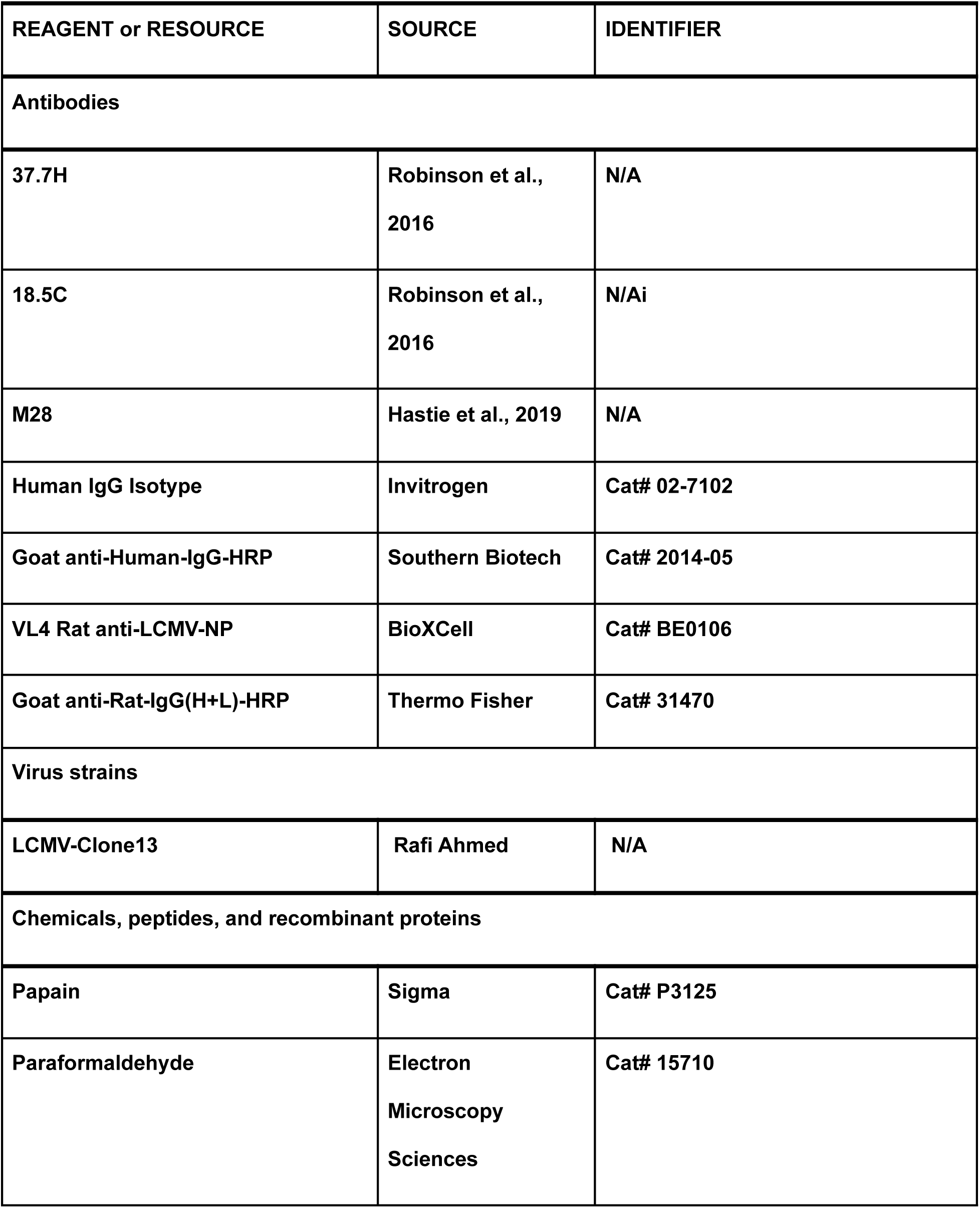
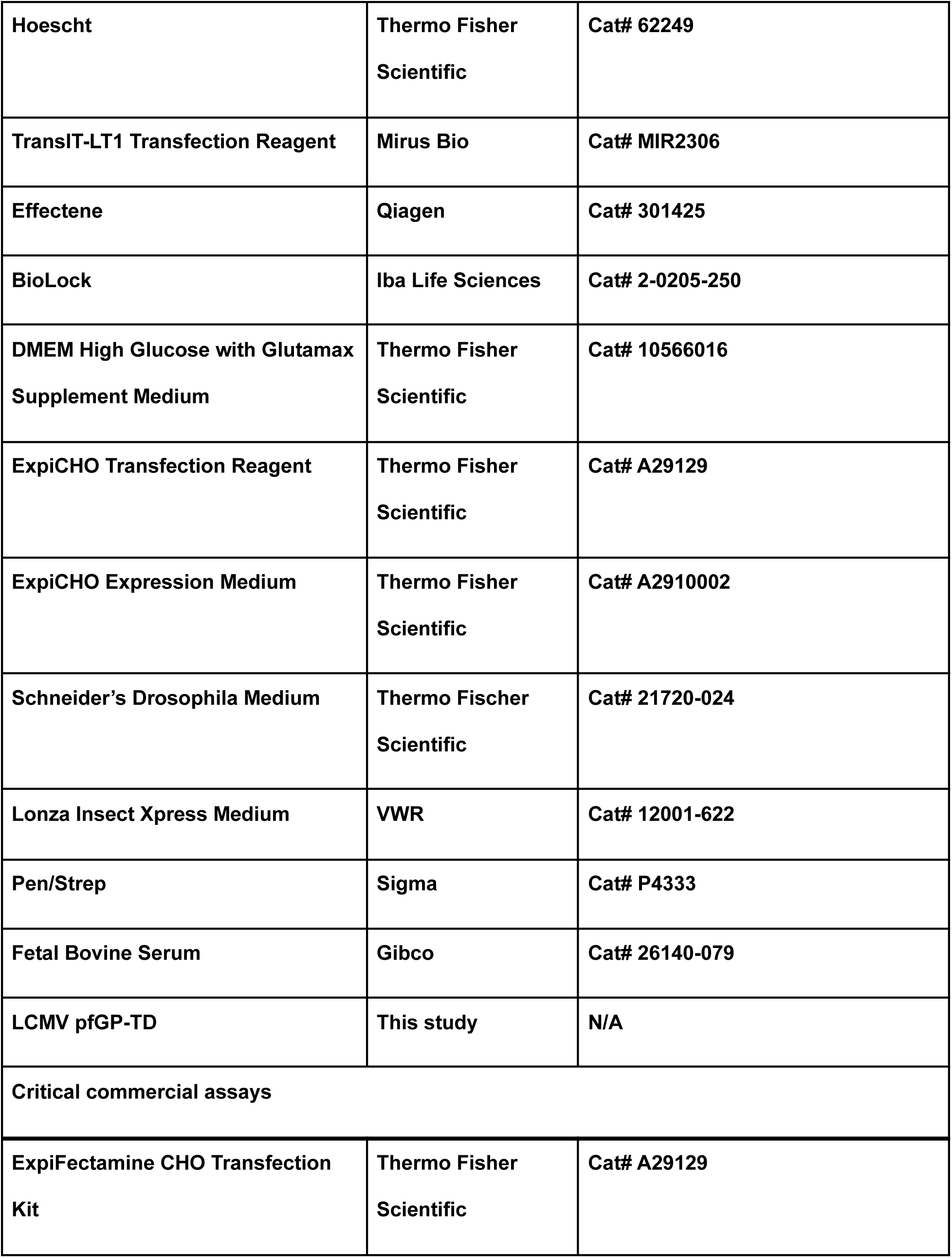
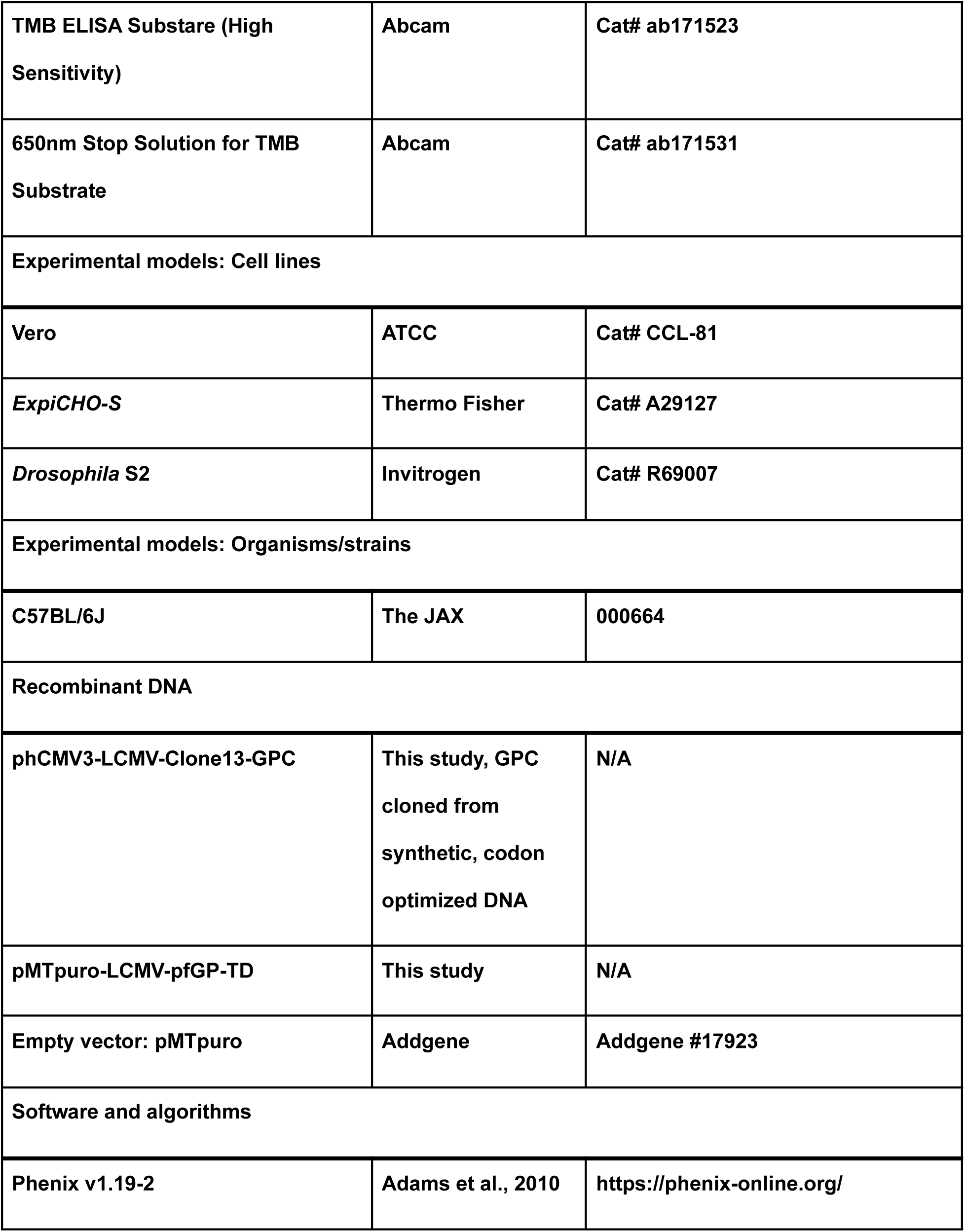
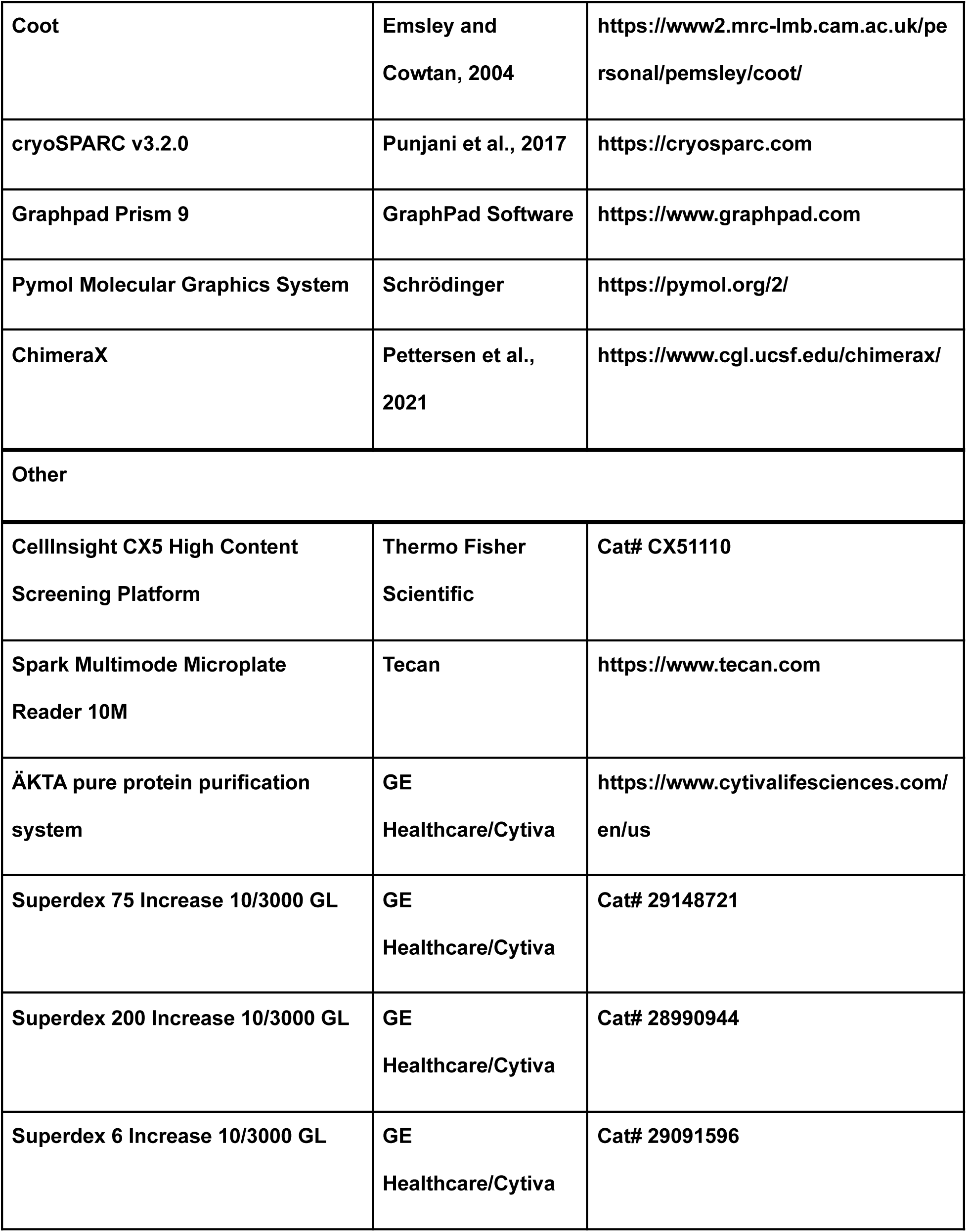
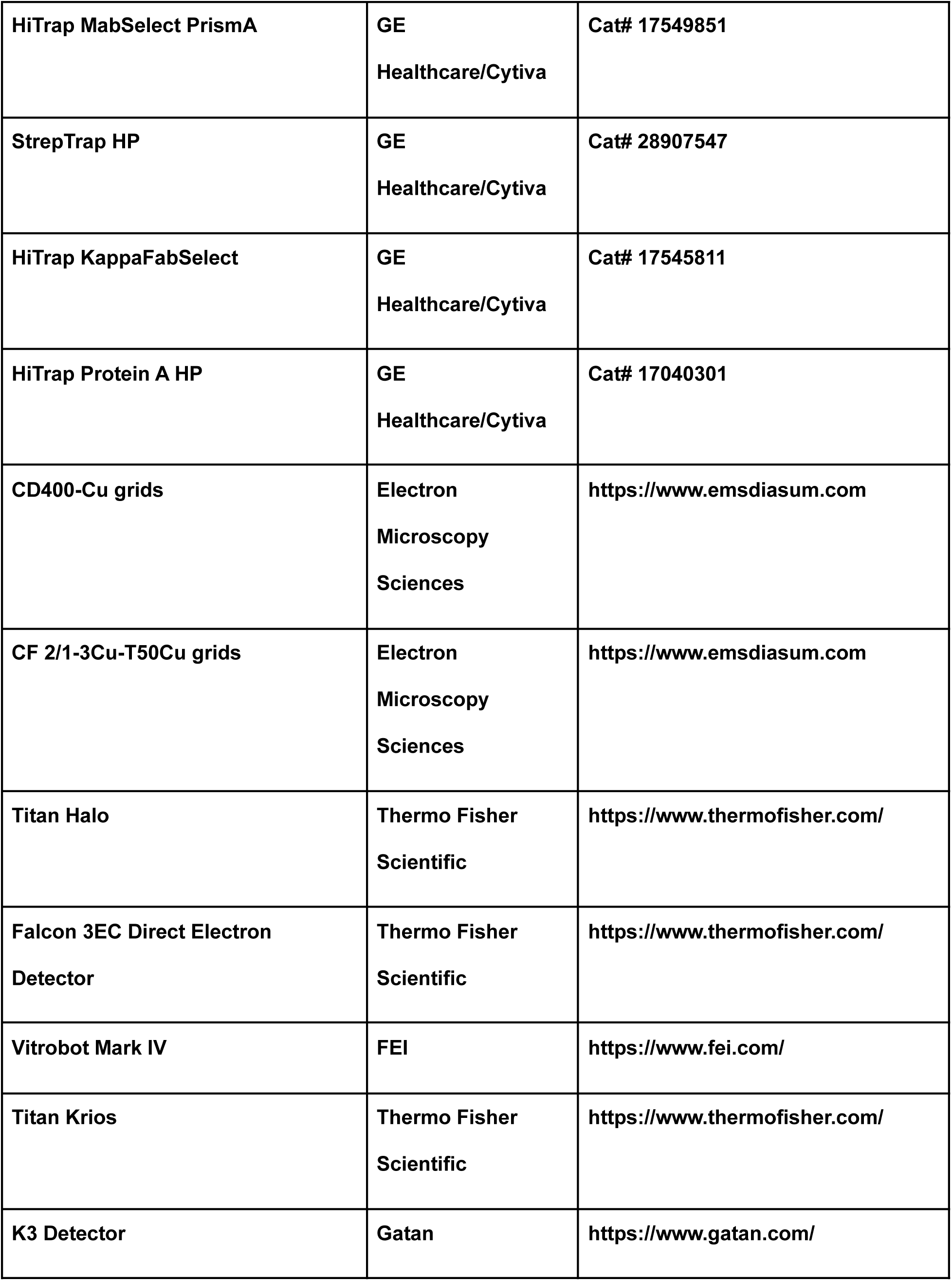

## Materials and Methods

### Cells

Vero cells (ATCC #CCL-81) were used for the LCMV neutralization and viral titer studies. These cell lines were cultured in Dulbecco’s modified Eagle’s medium (DMEM) supplemented with 10% fetal bovine serum (FBS) and penicillin–streptomycin (PS) and maintained at 37°C and 5% CO2. ExpiCHO cells (Thermo Fisher #A29127) were used in the production of IgG. *Drosophila* S2 cells (Invitrogen) were used to produce LCMV pfGP-TD.

### Mice, LCMV infection, and monoclonal antibody treatment

Mouse experiments were performed at La Jolla Institute for Immunology (LJI). All experimental procedures were approved by the IACUC committee of LJI. For prophylactic treatment, 6-weeks old male C57BL/6J mice were first injected with 150 µg M28 or IgG control via I.V. and 150 µg M28 or IgG control via I.P. (total is 300 µg) at day -1 and infected with medium-dose 0.3×10^6^ or high-dose 2×10^6^ FFU LCMV^cl13^ via I.V. at day 0. The blood or serum, spleen, one piece of liver, and kidneys were harvested on days 3 and 7, and the organs were weighed and homogenized in 1 ml DMEM with 2% FBS and PS (D2). For titrated infection of LCMV^cl13^, 6-weeks old male C57BL/6J mice were infected with 2×10^6^, 1×10^6^, or 0.3×10^6^ FFU LCMV^cl13^, serum was collected at day 7, day 15, and day 28 to determine viral titers. For therapeutic treatment, 6-weeks old male C57BL/6J mice were infected with 0.3×10^6^ FFU LCMV^cl13^ at day 0, 150 µg M28 or IgG control were injected via I.V. and I.P. (total is 300 µg) 2 days post-infection. Serum was collected on day 7, day 14, and day 21. Spleen, one piece of liver, and kidney were collected at day 21, weighed, and homogenized in D2. The weight of spleen and two kidneys from 10 weeks old mice were ∼0.075 g and ∼0.391g, respectively. The viral titers in organs were calculated as FFU/g. The viral titers in blood or serum were calculated as FFU/mL.

### Determination of LCMV titers

The LCMV titers were determined by immunofocus assay. In brief, 30 µl of blood, serum, or homogenized organ suspension were 10-fold serial diluted in D2 and mixed with 4×10^4^ Vero cells in 96-well plate at 37°C for 1 hour; then overlay (1% methylcellulose in D2) was added and incubated at 37°C for 2 days. On the third day, the supernatant was removed, the cells were fixed with 4% paraformaldehyde and permeabilized (0.1% Triton X-100 in PBS). After blocking (3% BSA in PBS), infectious foci were visualized using VL4 rat-anti-LCMV-NP antibody (BioXCell) and secondary HRP-conjugated goat-anti-rat-IgG (Thermo Fisher), followed by KPL-True Blue (Seracare). The reaction was stopped by washing with tap water. The plate images were captured by ELISPOT, and foci were counted from each well. The limited detection of virus in organs was 800 FFU/g in a 96-well plate. The limited detection of virus in serum is 50 FFU/mL. For immunofocus based neutralization assay, 2-fold serial diluted M28, 18.5C or 37.7H monoclonal antibody was mixed with 60 FFU LCMV^cl13^ at 37°C for 1.5 hours. Then the mixture was transferred onto the Vero cells, and the infectious foci were visualized as described above.

### Fab expression and purification

ExpiCHO cells were transfected with equimolar amounts of pLM-M28 light chain and pLM-M28 heavy chain expressing plasmids according to the manufacturer’s protocol (Thermo Fisher). At 12 days post-transfection, supernatant was cleared of cells via centrifugation. pH of the media was adjusted using a final concentration of 50mM Tris-HCl pH 8.0, followed by filtration using a 0.2μm filter. M28 IgG was then purified from the supernatant using Protein A affinity chromatography (GE Healthcare #17549851). IgG was then dialyzed overnight in TBS (150mM NaCl, 25mM Tris-HCl, pH 7.4). M28 Fab was obtained by cleaving M28 IgG with 5% papain (Sigma), followed by purification using a HiTrap KappaSelect column (GE Healthcare #17545811). Purified Fab was dialyzed in TBS (150mM NaCl, 25mM Tris-HCl, pH 7.4).

### Glycoprotein expression and purification

3×10^6 S2 cells (Invitrogen)/well were plated on a 6-well plate in Complete Schneider’s media supplemented with 10% FBS and pen/strep (CS10PS). The following day, cells were transfected with 2µg of pMT-PuroR-BiP-LCMV-pfGP-TD-EK-DS-Avi using Effectene Transfection Reagent (Qiagen) according to the manufacturer’s protocol. 18 hours post-transfection, cells were washed in PBS once using centrifugation. Washed cells were then added to a 6-well plate with fresh (CS10PS). 48 hours post transfection, puromycin was added to a final concentration of 6µg/mL. Puromycin resistant cells had media routinely replaced every 2 days and were allowed to expand to a T75 flask (Corning). Once confluent, cells were moved to a shaker flask with LONZA Insect Xpress serum free medium supplemented with 6 µg/mL puromycin. Cells were then expanded to a 2L final volume. Once cells reached a concentration of 1×10^7 cells/mL, protein expression was induced with 500µM CuSO_4_ and cells were allowed to produce protein for 4 days. After 4 days, supernatant was cleared from cells using centrifugation. The pH of supernatant was adjusted to pH 8.0 using 1M Tris-HCl and filtered using a 0.2µm filter to remove any cell debris. BioLock (IbaLife Sciences) was then added to the filtered supernatant according to the manufacturer’s protocol and allowed to incubate overnight at 4°C. The following day, the supernatant was filtered using a 0.2µm filter to remove any precipitates. Protein was then purified from the supernatant using a StrepTrap HP column (GE Healthcare #28907547), followed by SEC purification over a S6 increase column (GE Healthcare #29091596) in TBS (25mM Tris, 150mM NaCl, pH 7.4).

### ELISA

Half-area high protein-binding plates (Corning #07-200-37) were coated with 200ng/well of LCMV pfGP-TD in TBS (25mM Tris HCl, 150mM NaCl, pH 7.4) and incubated at 37 °C for 1 hour. Wells were then incubated with a blocking buffer composed of 3% BSA in TBS for 1 hour at 37 °C for 1 hour. Wells were washed three times in ELISA buffer (TBS supplemented with 0.1% Tween20 and 3% BSA). The indicated IgG resuspended in ELISA buffer in a 2-fold dilution series was then added to the plates and incubated and were incubated at 37 °C for 1 hour. Wells were washed six times in ELISA buffer and then incubated with 1:1000 dilution of goat α-human IgG-HRP in ELISA buffer 1 hour at 37°C. Wells were once again washed six times with ELISA buffer. TMB ELISA Substrate (Highest Sensitivity) (Abcam ab171522) was added, followed by equal volumes of 650 nm Stop Solution for TMB Substrate (Abcam ab171531). The OD650 of each well was read using a spectrophotomer plate reader (Tecan, Spark Multimode Reader).

### Negative Stain Electron Microscopy

SEC purified LCMV pfGP-TD-M28 Fab complex was diluted to a concentration of 0.03mg/mL in TBS (150mM NaCl, 25mM Tris-HCl, pH 7.4). 4µL of this diluted sample was applied to 15s 15mA glow discharge CD400-Cu negative stain grids (Electron Microscopy Sciences). Grids were stained using 0.8% uranyl formate. Micrographs were taken using a Titan Halo electron microscope (Thermo Fisher) and a Falcon 3EC direct electron detector at a 58,000X magnification. 2D classification and ab-initio reconstruction analysis was done using CryoSPARC^61^.

### CryoEM Sample Preparation

SEC-purified LCMV pfGP-TD and LCMV pfGP-TD/M28 Fab complex in TBS was concentrated to 0.8 and 1.2mg/mL using a 30K Amicon centrifugal filter (Sigma Aldrich). For LCMV pfGP-TDapo, 3µL of each sample was directly applied to a glow discharged C-Flat 2/1-3Cu-T50Cu grids (Electron Microscope Sciences). For the pfGP-TD-M28 complex, we used R2/2 grids (Quantifoil) coated with a thin layer of 0.2 mg/ml graphene oxide. Grids were equilibrated at 4°C in 100% humidity, followed by a 3-second blotting time using Vitrobot Mark IV (FEI) and then plunged into liquid ethane. Data were collected using the Titan Krios (Thermo Scientific) microscope operating at 300kV equipped with a K3 detector and BioQuantum energy filter with 20 eV slit (Gatan). A complete description of data collection parameters can be found in Table S1.

### CryoEM processing and atomic model building

Movies were processed using motion correction and CTF-estimation in CryoSPARC v3.2.0^61^. For both datasets (pfGP-TDapo and pfGP-TD/M28), particles from a subset of 2,000 micrographs were picked using Topaz^62^ in CryoSPARC and the output of this was used as templates for particle picking from the remaining micrographs. Extracted particles were subjected to reference-free 2D classification. Particles from selected 2D classes were then processed in RELION^63^. In RELION, particles underwent subsequent rounds of 2D classification, followed by 3D classification and 3D refinement. The finalized C3-symmetrized reconstructions for LCMV pf-GP-TDapo and LCMV pf-GP-TD-M28 reported a resolution of 3.26Å and 3.19Å. Final maps were processed in DeepEMhancer^64^. The model was built in Coot^65^ using the maps from DeepEMhancer and using the models of the uncleaved LCMV WE HPI strain GP_1-2_ (PDB 5INE)^45, 62^ and the parental 18.5C Fab structure (PBD 6P91)^39^ as templates. Refinement was carried out in PHENIX^66^ using the final 3D refinements models. Local resolution plots were calculated at FSC=0.5 in CryoSPARC. Figures were generated using ChimeraX^67^.

### Statistical analysis

Statistical analyses were performed in GraphPad Prism 9.3.0, unless otherwise stated. The statistical details of the experiments are provided in the respective figure legends. For statistical analysis of viral titers, two-tailed Student’s t-test or two-way ANOVA followed by Tukey’s post hoc test were performed by using log-transformed data. Statistical comparison of mice weight over 20 days was performed by two-way ANOVA.

## Notes

### Competing Interest Statement

The authors have declared no competing interest.

## References

1. Traub, E. THE EPIDEMIOLOGY OF LYMPHOCYTIC CHORIOMENINGITIS IN WHITE MICE. J. Exp. Med. 64, 183–200 (1936).

2. Park, J. Y. et al. Age distribution of lymphocytic choriomeningitis virus serum antibody in Birmingham, Alabama: evidence of a decreased risk of infection. Am. J. Trop. Med. Hyg. 57, 37–41 (1997).

3. Albariño, C. G. et al. High diversity and ancient common ancestry of lymphocytic choriomeningitis virus. Emerg. Infect. Dis. 16, 1093–1100 (2010).

4. Zapata, J. C. & Salvato, M. S. Arenavirus variations due to host-specific adaptation. Viruses 5, 241–278 (2013).

5. Zinkernagel, R. M. & Doherty, P. C. Immunological surveillance against altered self components by sensitised T lymphocytes in lymphocytic choriomeningitis. Nature 251, 547–548 (1974).

6. Moskophidis, D., Lechner, F., Pircher, H. & Zinkernagel, R. M. Virus persistence in acutely infected immunocompetent mice by exhaustion of antiviral cytotoxic effector T cells. Nature 362, 758–761 (1993).

7. Armstrong, C. & Lillie, R. D. Experimental Lymphocytic Choriomeningitis of Monkeys and Mice Produced by a Virus Encountered in Studies of the 1933 St. Louis Encephalitis Epidemic. Public Health Reports *(1896-1970)* vol. 49 1019 (1934).

8. Barton, L. L., Mets, M. B. & Beauchamp, C. L. Lymphocytic choriomeningitis virus: emerging fetal teratogen. Am. J. Obstet. Gynecol. 187, 1715–1716 (2002).

9. Fischer, S. A. et al. Transmission of lymphocytic choriomeningitis virus by organ transplantation. N. Engl. J. Med. 354, 2235–2249 (2006).

10. Jamieson, D. J., Kourtis, A. P., Bell, M. & Rasmussen, S. A. Lymphocytic choriomeningitis virus: an emerging obstetric pathogen? Am. J. Obstet. Gynecol. 194, 1532–1536 (2006).

11. Delaine, M. et al. Microcephaly Caused by Lymphocytic Choriomeningitis Virus. Emerg. Infect. Dis. 23, 1548–1550 (2017).

12. Kinori, M., Schwartzstein, H., Zeid, J. L., Kurup, S. P. & Mets, M. B. Congenital lymphocytic choriomeningitis virus-an underdiagnosed fetal teratogen. J. AAPOS 22, 79–81.e1 (2018).

13. Gregg, M. B. Recent outbreaks of lymphocytic choriomeningitis in the United States of America. Bull. World Health Organ. 52, 549–553 (1975).

14. Dräger, S. et al. Lymphocytic choriomeningitis virus meningitis after needlestick injury: a case report. Antimicrob. Resist. Infect. Control 8, 77 (2019).

15. Aebischer, O., Meylan, P., Kunz, S. & Lazor-Blanchet, C. Lymphocytic choriomeningitis virus infection induced by percutaneous exposure. Occup. Med. 66, 171–173 (2016).

16. Froeschke, M., Basler, M., Groettrup, M. & Dobberstein, B. Long-lived signal peptide of lymphocytic choriomeningitis virus glycoprotein pGP-C. J. Biol. Chem. 278, 41914–41920 (2003).

17. York, J. & Nunberg, J. H. Intersubunit interactions modulate pH-induced activation of membrane fusion by the Junin virus envelope glycoprotein GPC. J. Virol. 83, 4121–4126 (2009).

18. Shankar, S. et al. Small-Molecule Fusion Inhibitors Bind the pH-Sensing Stable Signal Peptide-GP2 Subunit Interface of the Lassa Virus Envelope Glycoprotein. J. Virol. 90, 6799–6807 (2016).

19. Burri, D. J. et al. The role of proteolytic processing and the stable signal peptide in expression of the Old World arenavirus envelope glycoprotein ectodomain. Virology 436, 127–133 (2013).

20. Kunz, S., Edelmann, K. H., de la Torre, J.-C., Gorney, R. & Oldstone, M. B. A. Mechanisms for lymphocytic choriomeningitis virus glycoprotein cleavage, transport, and incorporation into virions. Virology 314, 168–178 (2003).

21. Beyer, W. R., Pöpplau, D., Garten, W., von Laer, D. & Lenz, O. Endoproteolytic processing of the lymphocytic choriomeningitis virus glycoprotein by the subtilase SKI-1/S1P. J. Virol. 77, 2866–2872 (2003).

22. Buchmeier, M. J., Southern, P. J., Parekh, B. S., Wooddell, M. K. & Oldstone, M. B. Site-specific antibodies define a cleavage site conserved among arenavirus GP-C glycoproteins. J. Virol. 61, 982–985 (1987).

23. Recher, M. et al. Deliberate removal of T cell help improves virus-neutralizing antibody production. Nat. Immunol. 5, 934–942 (2004).

24. Cao, W. et al. Identification of alpha-dystroglycan as a receptor for lymphocytic choriomeningitis virus and Lassa fever virus. Science 282, 2079–2081 (1998).

25. Kunz, S., Sevilla, N., McGavern, D. B., Campbell, K. P. & Oldstone, M. B. Molecular analysis of the interaction of LCMV with its cellular receptor [alpha]-dystroglycan. J. Cell Biol. 155, 301–310 (2001).

26. Kanagawa, M. et al. Molecular Recognition by LARGE Is Essential for Expression of Functional Dystroglycan. Cell vol. 117 953–964 (2004).

27. Kunz, S. et al. Posttranslational modification of alpha-dystroglycan, the cellular receptor for arenaviruses, by the glycosyltransferase LARGE is critical for virus binding. J. Virol. 79, 14282–14296 (2005).

28. Volland, A. et al. Heparan sulfate proteoglycans serve as alternative receptors for low affinity LCMV variants. PLoS Pathog. 17, e1009996 (2021).

29. Bakkers, M. J. G. et al. CD164 is a host factor for lymphocytic choriomeningitis virus entry. Proc. Natl. Acad. Sci. U. S. A. 119, e2119676119 (2022).

30. Borrow, P. & Oldstone, M. B. Mechanism of lymphocytic choriomeningitis virus entry into cells. Virology 198, 1–9 (1994).

31. Di Simone, C., Zandonatti, M. A. & Buchmeier, M. J. Acidic pH triggers LCMV membrane fusion activity and conformational change in the glycoprotein spike. Virology 198, 455–465 (1994).

32. Eschli, B. et al. Early antibodies specific for the neutralizing epitope on the receptor binding subunit of the lymphocytic choriomeningitis virus glycoprotein fail to neutralize the virus. J. Virol. 81, 11650–11657 (2007).

33. Richter, K. & Oxenius, A. Non-neutralizing antibodies protect from chronic LCMV infection independently of activating FcγR or complement. Eur. J. Immunol. 43, 2349–2360 (2013).

34. Stoycheva, D. et al. Non-neutralizing antibodies protect against chronic LCMV infection by promoting infection of inflammatory monocytes in mice. Eur. J. Immunol. 51, 1423–1435 (2021).

35. Hangartner, L. et al. Nonneutralizing antibodies binding to the surface glycoprotein of lymphocytic choriomeningitis virus reduce early virus spread. J. Exp. Med. 203, 2033–2042 (2006).

36. Seiler, P. et al. Enhanced virus clearance by early inducible lymphocytic choriomeningitis virus-neutralizing antibodies in immunoglobulin-transgenic mice. J. Virol. 72, 2253–2258 (1998).

37. Fallet, B. et al. Chronic Viral Infection Promotes Efficient Germinal Center B Cell Responses. Cell Rep. 30, 1013–1026.e7 (2020).

38. Robinson, J. E. et al. Most neutralizing human monoclonal antibodies target novel epitopes requiring both Lassa virus glycoprotein subunits. Nat. Commun. 7, 11544 (2016).

39. Hastie, K. M. et al. Convergent Structures Illuminate Features for Germline Antibody Binding and Pan-Lassa Virus Neutralization. Cell 178, 1004–1015.e14 (2019).

40. Hastie, K. M. et al. Structural basis for antibody-mediated neutralization of Lassa virus. Science 356, 923–928 (2017).

41. Richmond, J. K. & Baglole, D. J. Lassa fever: epidemiology, clinical features, and social consequences. BMJ 327, 1271–1275 (2003).

42. McCormick, J. B. & Fisher-Hoch, S. P. Lassa fever. Curr. Top. Microbiol. Immunol. 262, 75–109 (2002).

43. Buck, T. K. et al. Neutralizing Antibodies against Lassa Virus Lineage I. MBio e0127822 (2022).

44. Enriquez, A. S. et al. Delineating the mechanism of anti-Lassa virus GPC-A neutralizing antibodies. Cell Rep. 39, 110841 (2022).

45. Hastie, K. M. et al. Crystal structure of the prefusion surface glycoprotein of the prototypic arenavirus LCMV. Nat. Struct. Mol. Biol. 23, 513–521 (2016).

46. Katz, M. et al. Structure and receptor recognition by the Lassa virus spike complex. Nature (2022) doi:10.1038/s41586-022-04429-2.

47. Li, S. et al. Acidic pH-Induced Conformations and LAMP1 Binding of the Lassa Virus Glycoprotein Spike. PLoS Pathog. 12, e1005418 (2016).

48. Saridakis, V. et al. The structural basis for methylmalonic aciduria. The crystal structure of archaeal ATP:cobalamin adenosyltransferase. J. Biol. Chem. 279, 23646–23653 (2004).

49. Dylla, D. E., Xie, L., Michele, D. E., Kunz, S. & McCray, P. B., Jr. Altering α-dystroglycan receptor affinity of LCMV pseudotyped lentivirus yields unique cell and tissue tropism. Genet. Vaccines Ther. 9, 8 (2011).

50. Kunz, S., Sevilla, N., Rojek, J. M. & Oldstone, M. B. A. Use of alternative receptors different than alpha-dystroglycan by selected isolates of lymphocytic choriomeningitis virus. Virology 325, 432–445 (2004).

51. Kunz, S., Rojek, J. M., Perez, M., Spiropoulou, C. F. & Oldstone, M. B. A. Characterization of the interaction of lassa fever virus with its cellular receptor alpha-dystroglycan. J. Virol. 79, 5979–5987 (2005).

52. Acciani, M. et al. Mutational Analysis of Lassa Virus Glycoprotein Highlights Regions Required for Alpha-Dystroglycan Utilization. J. Virol. 91, (2017).

53. Ishii, A. et al. Molecular surveillance and phylogenetic analysis of Old World arenaviruses in Zambia. J. Gen. Virol. 93, 2247–2251 (2012).

54. Palacios, G. et al. A New Arenavirus in a Cluster of Fatal Transplant-Associated Diseases. New England Journal of Medicine vol. 358 991–998 (2008).

55. Harker, J. A., Lewis, G. M., Mack, L. & Zuniga, E. I. Late interleukin-6 escalates T follicular helper cell responses and controls a chronic viral infection. Science 334, 825–829 (2011).

56. Greczmiel, U., et al. Sustained T follicular helper cell response is essential for control of chronic viral infection. Sci Immunol 2, (2017).

57. De Giovanni, M. et al. Spatiotemporal regulation of type I interferon expression determines the antiviral polarization of CD4 T cells. Nat. Immunol. 21, 321–330 (2020).

58. Sommerstein, R. et al. Arenavirus Glycan Shield Promotes Neutralizing Antibody Evasion and Protracted Infection. PLoS Pathog. 11, e1005276 (2015).

59. Tanveer, F., Younas, M. & Fishbain, J. Lymphocytic choriomeningitis virus meningoencephalitis in a renal transplant recipient following exposure to mice. Transpl. Infect. Dis. 20, e13013 (2018).

60. Mathur, G. et al. High clinical suspicion of donor-derived disease leads to timely recognition and early intervention to treat solid organ transplant-transmitted lymphocytic choriomeningitis virus. Transpl. Infect. Dis. 19, (2017).

61. Punjani, A., Rubinstein, J. L., Fleet, D. J. & Brubaker, M. A. cryoSPARC: algorithms for rapid unsupervised cryo-EM structure determination. Nat. Methods 14, 290–296 (2017).

62. Bepler, T., Kelley, K., Noble, A. J. & Berger, B. Topaz-Denoise: general deep denoising models for cryoEM and cryoET. Nat. Commun. 11, 5208 (2020).

63. Scheres, S. H. W. RELION: implementation of a Bayesian approach to cryo-EM structure determination. J. Struct. Biol. 180, 519–530 (2012).

64. Sanchez-Garcia, R. et al. DeepEMhancer: a deep learning solution for cryo-EM volume post-processing. Commun Biol 4, 874 (2021).

65. Emsley, P. & Cowtan, K. Coot: model-building tools for molecular graphics. Acta Crystallogr. D Biol. Crystallogr. 60, 2126–2132 (2004).

66. Adams, P. D. et al. PHENIX: building new software for automated crystallographic structure determination. Acta Crystallogr. D Biol. Crystallogr. 58, 1948–1954 (2002).

67. Pettersen, E. F. et al. UCSF ChimeraX: Structure visualization for researchers, educators, and developers. Protein Sci. 30, 70–82 (2021).

